# C3: Connect separate Connected Components to form a succinct disease module

**DOI:** 10.1101/2020.06.05.137398

**Authors:** Bingbo Wang, Jie Hu, Chenxing Zhang, Yuanjun Zhou, Liang Yu, Xingli Guo, Lin Gao, Yunru Chen

## Abstract

Accurate disease module is helpful in understanding the molecular mechanism of disease causation and identifying drug target. However, for the fragmentization of disease module in incomplete human interactome, how to determine connectivity pattern and detect a full neighbourhood of disease is an open problem. In this paper, a topology-based method is developed to dissect the connectivity of intermediate nodes and edges and form a succinct disease module. By applying this Connect separate Connected Components (CCC, C3) method on a large corpus of curated diseases, we find that most Separate Connected Components (SCCs) formed by Disease-Associated Proteins (DAPs) can be connected into a well-connected component as a succinct observable module. This pattern also holds for altered genes from multi-omics data such as The Cancer Genome Atlas. Overall, C3 tool can not only inspire a deeper understanding of interconnectedness of phenotypically related genes and different omics data, but also be used to detect a well-defined neighbourhood that drives complex pathological processes.

## Introduction

Most complex human diseases are rarely the consequence of the abnormal activity of a single gene product. Rather, they involve the activities of multiple interactional gene products, many of which otherwise carry no defect^1^. Hence, it is crucial to study genotype - phenotype relationships in the context of human interactome (Molecular interactions network, nodes are proteins and edges are built on observational inference of their interactions), to disentangle the genetic origin of complex diseases ^2–6^.

Previous work, driven by high-throughput interactome mapping efforts^7^ and the wealth of genome-wide genetic association data^8^, has begun to unveil the relationships between distinct phenotypes based on network. The most important concept to emerge from this is that of the “disease module” - a well-defined neighbourhood of **D**isease-**A**ssociated **P**roteins, DAPs (encoded by disease genes) clustered in the network that drives specific disease processes^3^. Topological properties of disease module in the interactome underlie all network-based prioritization tools for studying human diseases. DAPs implicated in a specific phenotype frequently form a well-connected component^9^ rather than locally dense community. And the vast majority of disease genes show no tendency to encode hub proteins and localize in the functional periphery of interactome^1^. Furthermore, the location of each disease module related to others in the interactome implies its phenotypic similarity to other diseases^10^. And recently, Jure Leskovec, et al^11^ have found that DAPs of a single disease tend to form many separate connected components in the network. They also showed that higher-order network structures, such as small subgraphs of the pathway, provide a promising direction for the development of new methods.

Disease module helps uncover the molecular mechanism of disease causation, identify new disease genes and pathways and aid rational drug target identification^9,12–18^. For example, Amitabh Sharma et al^18^ identified the asthma disease module and validated its functional and pathophysiological relevance, using both computational and experimental approaches. They found that the asthma disease module is enriched with modest GWAS p-values against the background of random variation, and with differentially expressed genes from normal and asthmatic fibroblast cells treated with an asthma-specific drug. The asthma module also contains immune response mechanisms that are shared by other immune-related disease modules.

Numerous computational methods have been developed for accurate identification of disease module to fully explain the molecular mechanism of most human diseases. For the fragmentation of disease module^10^, over the respective 80% DAPs appear to be distributed on average, forming many **S**eparate **C**onnected **C**omponents (SCCs), this pattern holds for more than 90% diseases^11^. A disease module does not correspond to a single well-connected component as an observable module in present incomplete interactome. Therefore, methods^9,14,19,20^ used for searching for dense clusters or communities of DAPs in the interactome, might be limited in discovering disease module. Such as the state-of-the-art algorithm DIAMOnD^9^ presented an analysis of the connectivity patterns of DAPs and formed a disease module with a large number of intermediate proteins, which are called DIAMOnD proteins. Although DIAMOnD can generate an interconnected module for a specific disease, it unfavourably imports 200 DIAMOnD proteins to connect only about average 50% DAPs from Online Mendelian Inheritance in Man (OMIM^21^) and Genome-Wide Association Studies (GWAS^22^) databases (Figure 1A). Many of these imported DIAMOnD proteins are unnecessary for connectivity, resulting in enormous sizes (average size 400 proteins, that is, the disease module has been amplified 4 times) but ambiguous disease associations of final disease modules in 299 disease phenotypes. Another kind of algorithms is based on Steiner tree, retrieval of a minimum size sub-graph leads to the classical Steiner tree problem, which is known as NP-complete^23^, such as a heuristic approach minimum spanning tree based approximation, the whole algorithm has time complexity about O(n^3^). It is unpractical due to an time-consuming hunt for optimal connected component in the huge interactome with more than 13,000 proteins. In addition, some remote nodes can also be connected by a Steiner tree, connectivity significance of disease module thus be hindered by such issues, may lead to misunderstanding of the topological characteristics of DAPs and disease modules. Tumminello, M., et al introduce a method^24^ to detect network structures. They associate a p-value with each edge of the network in order to test the presence of the edge against the selected null hypothesis, and generate statistically validated networks. Percolation in physics can also be understood in a broad sense as processes related to the connection of nodes of a network, and is generally associated with the emergence of a giant cluster at a given connection probability^25^. Furthermore, although numerous methods exist, analysis of connectivity patterns and detection statistically significant of DAPs remains an open problem.

**Figure 1.**
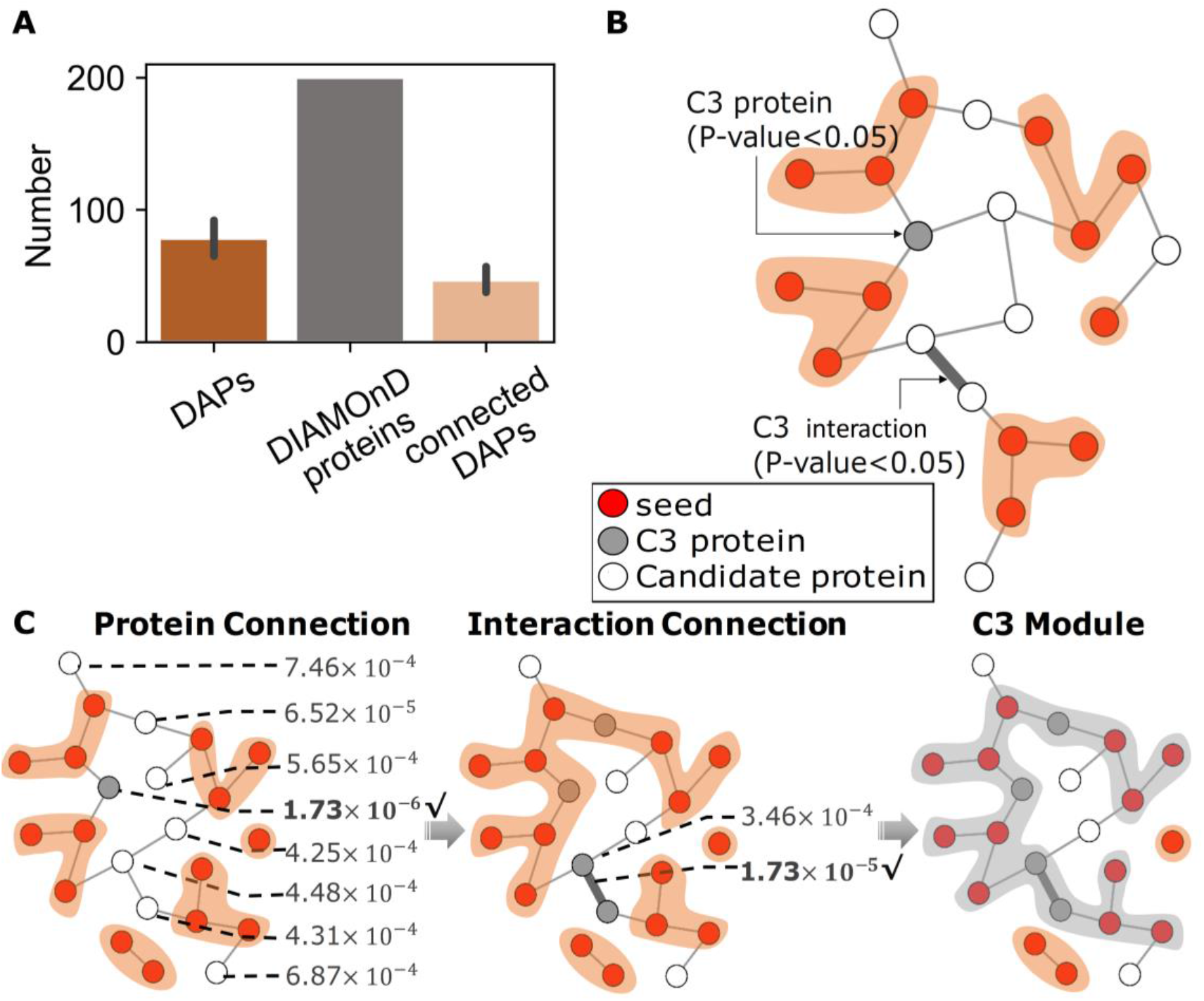
A schematic diagram of C3 module. (A) The redundancy of disease modules of the 299 diseases obtained from the DIAMOnD. The dark orange bars represent the average number (about 78) of seeds with 299 diseases, the gray bars indicate that 200 DIAMOnD proteins are obtained for each disease using the DIAMOnD, and the light orange bars indicate that these intermediates only connected about 50% seeds in final disease module. The error bar indicates a 95% confidence interval. (B) The definition of a C3 protein and a C3 interaction. The yellow shade is SCCs. Connectivity of C3 protein and C3 interaction are quantified by hypergeometric test P-values with threshold p-values < 0.05. (C) Schematic diagram of the C3 method. At each step, we regard immediate neighbors of seeds as candidate proteins, first we carry out protein connection operation namely calculate the connectivity probability (p-value) of all candidate proteins. Then, the protein with minimum p-value is selected as a C3 protein to connect SCCs and is considered as a seed in the next step. Next, if it can’t connect at least two SCCs, we carry out interaction connection operation, do the same for the edge of candidate proteins participated, the proteins occupied the endpoints of a C3 interaction are also considered as seeds in the next step. Thus, after several steps, when interaction connection operation also can’t connect at least two SCCs, we obtain the C3 module (gray shadow).

Here we develop a topology-based method to **C**onnect separate **C**onnected **C**omponents (CCC, C3) to form a succinct disease module. It is established that SCCs of DAPs can be connected effectively through a few intermediate proteins and interactions in the network, which can significantly reduce the number of fragments of a disease module in statistics. First, we quantify the roles of proteins and interactions in connecting SCCs. By importing the intermediate proteins, C3 indicates observable disease modules of DAPs from OMIM and GWAS databases for 299 disease phenotypes. Then we describe the characteristics of our disease modules, we validate the succinctness and connectivity significance of C3 module by comparing the number of imported intermediate proteins with results of DIAMOnD and random simulations. A well - connected succinct disease module refers to one small size and dominated by seed genes. Furthermore, we explore their relative contributions to disease pathogenesis in a few case studies. And we present evidence that even for DAPs from multi-omics data in The Cancer Genome Atlas (TCGA^26^), C3 works as a practical strategy for understanding interconnectedness of mRNA expression, mutation, DNA methylation and copy number variation, further validating the biological significance of the disease module we obtained.

## Results

### Connectivity pattern of disease modules generated by C3

We obtain DAPs by mapping disease genes into the human interactome. We call them “seeds”. To identify disease module and explore its connectivity pattern, we start by quantifying the abilities of intermediate proteins or interactions to connect SCCs of seeds. For each disease, we project its seeds (seeds set *D*) into the PPI network *G*(*N*, *E*), where *N* is the set of proteins and *E* is the set of interactions, then obtain the SCCs (connected components set *S*). For a protein *i* ∈ *N* or an interaction *e* ∈ *E* in the immediate neighbours of *D*, the hypergeometric p-value (see Methods) is used to indicate the significance of a protein *i* or an interaction *e* in connecting different elements in set *S*. We define a *C3 protein* or a *C3 interaction* as a protein or an interaction that can significantly connect SCCs (with p-value < 0.05, that is connectivity is significant) in statistics (Figure 1B). The C3 proteins and the proteins occupied the endpoints of C3 interactions work as intermediates, which are uniformly referred to as “*C3 proteins*” in the following functional analysis. Then, by a greedy procedure (see Methods), importing C3 proteins and C3 interactions, when there is no C3 proteins and C3 interactions that can connect SCCs with p-value < 0.05, the process is stopped, a well-connected disease module (named *C3 module*) becomes observable (Figure 1C). Furthermore, the universality of C3 proteins and C3 interactions for various diseases provides further evidences for the connectivity pattern that DAPs intend to form a well-defined and detectable neighbourhood in the human interactome.

Next, C3 method is applied to exhibit the disease modules in a comprehensive list of experimentally documented molecular interactions in human cells compiled by Menche, J., et al^10^ (see Methods). In total, we obtain 141,296 interactions between 13,460 proteins, 1,531 of which are associated with one or more diseases. The known disease genes are gathered from OMIM^21^ and GWAS^22^ databases (see Methods). Then the connectivity patterns of 299 well-characterized complex diseases in a curated list are studied (see results in supplementary files). The results of these various diseases confirm the universality of C3 proteins and C3 interactions that can be used as intermediates for disease modules. Furthermore, to validate more general applicability of C3 method, we collect multi-omics data including mRNA expression, mutations, DNA methylation and copy number variations from TCGA (see Methods), and successfully identify multi-omics neighbourhoods of Breast Cancer (BRCA).

### Succinctness and connectivity significance of the C3 module

According to disease module hypothesis, genes associated with a specific disease phenotype show a tendency to cluster in the same network neighbourhood, often referred to as a “disease module”3. Furthermore, the location of each disease module relative to others in the network implies its phenotypic similarity to other diseases^10^, it is important to position the disease module in the network. In DIAMOnD module, due to the large number of the DIAMOnD proteins, DIAMOnD disease module is a huge one dominated by DIAMOnD proteins, it will bring many false signals to the disease module. This is why we form a succinct disease module. Since the C3 disease module imports a small amount of C3 proteins and is dominated by DAPs, the C3 disease module can be accurately located on the network and provide us with more accurate disease signals. So a succinct disease module is essential for studying disease occurrence and treatment. The succinctness of the disease module means that for DAPs, a disease module can be constructed with as few intermediate proteins as possible.

As is mentioned above, disease modules of the 299 diseases obtained from the DIAMOnD algorithm are redundant. More favorable for DIMAMOnD, we compare C3 with DIAMOnD algorithm on a same data platform, the supporting data of DIAMOnD includes: human interactome network and the curated list of 299 diseases. In addition, we use current or systematically generated networks BioGRID, Bioplex and HuRI. Network information is in Table 1. In human interactome, by observing the fraction of connected seeds when they use the same intermediate proteins, we observe that C3 connected seeds are about 95%, and DIAMOnD is about 39%, the p-value of their difference is 3.94e-99. The other three networks are the same, P-values are 5.80e-98, 4.09e-96, 1.58e-57 respectively (Figure 2A). In the end, it is validated that when importing the same intermediate proteins, C3 can connect more seeds efficiently than DIAMOnD, so we can find the succinctness of the C3 module.

**Table 1.** Network Information.

| Network | Source | #nodes<br>#edges | Mean<br>degree | Diameter | Clustering<br>coefficient |
| --- | --- | --- | --- | --- | --- |
| Human<br>Interactome | Menche, J. et al. 2015 | 13,460<br>141,296 | 20.6 | 13 | 0.17 |
| BioGRID | https://thebiogrid.org/ | 17,741<br>338,793 | 37.9 | 8 | 0.12 |
| Bioplex | https://bioplex.hms.harvard.edu/ | 10,961<br>56,553 | 10.3 | 11 | 0.09 |
| HuRI | http://interactome.dfci.harvard.edu/H_sapiens/ | 7,352<br>38,414 | 10.3 | 11 | 0.06 |

**Figure 2.**
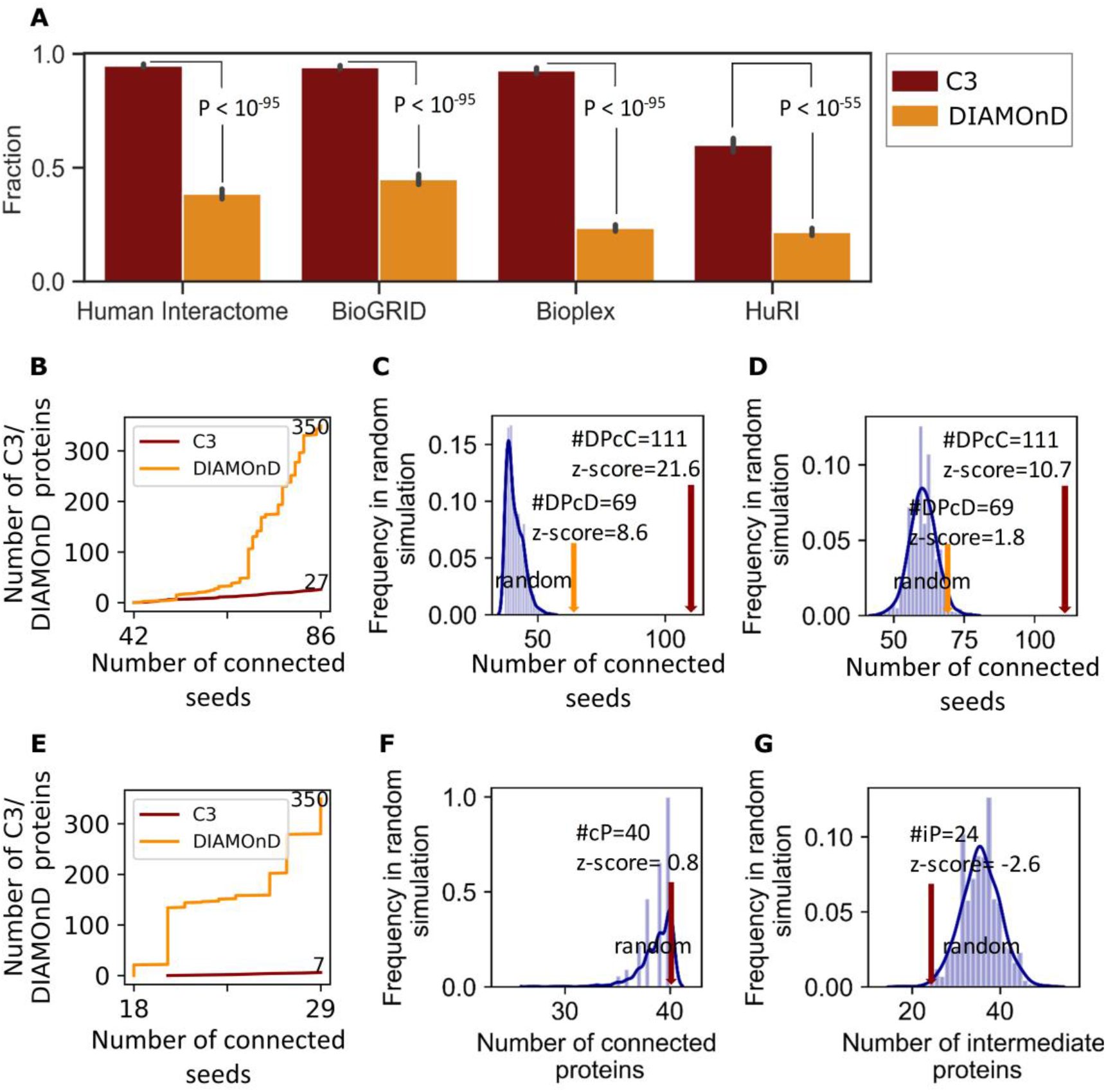
Succinctness and connectivity significance of the C3 module. (A) The fraction of seeds contained in the C3 and DIAMOnD modules of 299 diseases in different networks, and p-value obtain by wilcoxon test. (B, C, D) For asthma, (B) when the modules obtained by DIAMOnD and C3 contain 86 seeds at the same time, red line indicates that only 27 C3 proteins are used in the C3 module, while yellow line indicates that the DIAMOnD module used 350 DIAMOnD proteins. (C) Comparison of the number of Disease Proteins contained in DIAMOnD module (DPcD) and C3 module (DPcC) with 1000 random simulations. Modules constructed completely randomly contain on average 44.9 ± 3.2 seeds, which is significantly lower than the value in the real disease module. And DIAMOnD module contains 69 seeds (z-score = 8.6) lower than C3 module with 111 seeds (z-score = 21.6). (D) The similar as B, but choosing the intermediate proteins not completely at random, but only from the immediate neighbor of seeds. The random modules contain 60.3 ± 4.7 seeds, again significantly lower than in the real disease module. And DIAMOnD module contains 69 seeds (z-score = 1.8) also lower than C3 module with 111 seeds (z-score = 10.7). (E, F, G) With BRCA have 40 DAPs (seeds), (E) Same as B, when DIAMOnD and C3 module contain 29 seeds at the same time, only 7 C3 proteins are used in the C3 module while the DIAMOnD module used 350 DIAMOnD proteins. Comparison of the number of connected Proteins (cP) (F) and the number of intermediate Proteins (iP) (G) in the final C3 module with the number obtained in 1000 random simulations. Randomly select any 40 proteins with the same degrees as seeds each time. The number of connected seeds is on average 38.7 ± 1.6, where all 40 BRCA proteins connected by C3. The number of intermediate proteins used on average is 35.6 ± 4.4, where only 24 C3 proteins are used in BRCA C3 module.

Then we choose asthma and BRCA for detailed analysis (Figure 2B-2G). For asthma, the same 86 asthma seeds are connected by the DIAMOnD and C3, C3 proteins require 27 but DIAMOnD uses 350 intermediate proteins, which are about 13 times that of C3. The same for BRCA, we connect 29 BRCA seeds with 7 C3 proteins and 350 DIAMOnD proteins, which are about 50 times that of C3 (Figure 2B and 2E). So, using less C3 proteins can connect the same number of the seeds, further prove the succinctness of C3 method. C3 method is equally applicable to other diseases. The succinctness of the C3 module is manifested in two ways, for connecting the same number of seeds, less C3 proteins are used. In addition, if the same number of intermediate proteins is used, C3 proteins can connect more seeds.

Connectivity significance refers to the universal presence of C3 proteins and C3 interactions in the network. They can significantly connect SCCs of seeds, that is, can play a role in the connectivity of disease module. To test whether the observed disease modules represent non-random aggregation of DAPs, we do random experiments on asthma and BRCA. For asthma, we find that modules of the same size (add 100 intermediate proteins as DIAMOnD) that are constructed completely at random on average contain 44.9 ± 3.2 seeds, which is significantly lower than in the real disease module, both DIAMOnD module (contain 69 seeds, z-score = 8.6) and C3 module (contain 111 seeds only used 61 C3 proteins, z-score = 21.6) (Figure 2C), even when the random proteins are chosen only from the immediate neighbors of seeds, we find that the resulting modules contain only 60.3 ± 4.7 seeds, which is again significantly lower as observed in the DIAMOnD module (z-score = 1.8) and C3 module (z-score = 10.7) (Figure 2D). For BRCA with 40 DAPs, we randomly select 40 proteins with the same degrees as seeds for random experiments. In the result we use 35.6 ± 4.4 intermediate proteins to connect 38.7 ± 1.6 seeds which are randomly selected. It can be seen that C3 not only connects all 40 seeds (z-score = 0.8) and uses significantly less intermediate proteins (z-score = −2.6) compared to random simulations (Figure 2F and 2G). These differences suggest that C3 successfully identifies connectivity significance of the disease module in the interactome, indicating the significant tendency of DAPs aggregation. And the C3 disease module outperforms DIAMOnD in revealing the connected pattern of DAPs in the interactome.

### C3 module in asthma

We have acquired C3 modules of 299 diseases. Whether can these modules effectively characterize the corresponding diseases? In the following, we use asthma C3 module for analysis and comparision with DIAMOnD module. The asthma test mentioned above is based on 139 DAPs from five sources: (i) Literature; (ii) GWAS; (iii) OMIM; (iv) MeSH; (v) Pathways^18^, 129 (92.8%) of which are represented in the human interactome network. Then we project them onto the interactome and obtain the SCCs (Figure 3A). It was observed that the seeds form a huge number of SCCs (83 SCCs) in the real interactome. Then, C3 method is used to connect these SCCs and obtain a well-connected module of asthma. We use 61 C3 proteins to well connect 65 (78%) SCCs, inferring that the SCCs tend to be interconnected rather than dispersed (Figure 3B). In addition, we need bioinformatics to verify the significance of this asthma C3 module. For unconfirmed C3 proteins in this module, the associations between them and asthma have not yet been discovered, and we have hinted at this possibility through enrichment analysis, which provides clues for the diagnosis and treatment of asthma. We use three different asthma-specific validation data: (i) Gene expression of asthma specific compiled from eight sources; (ii) Gene Ontology (GO) term similarity (Biological Processes, BP); (iii) Disgenet (see Materials and Methods and Figure 3C). The GO term similarity is calculated by using the tool GOSemSim^28^. In all three assessments, we observe that enrichment number of C3 and DIAMOnD proteins can reach the same level. And the level of enrichment p-value of C3 is as significant as DIAMOnD and even better sometimes. These three enrichment analyses show that C3 proteins have biological significance, inferring it is biologically related to asthma, so the 61 C3 proteins, together with the connected seeds which form C3 module, represent a credible asthma module.

**Figure 3.**
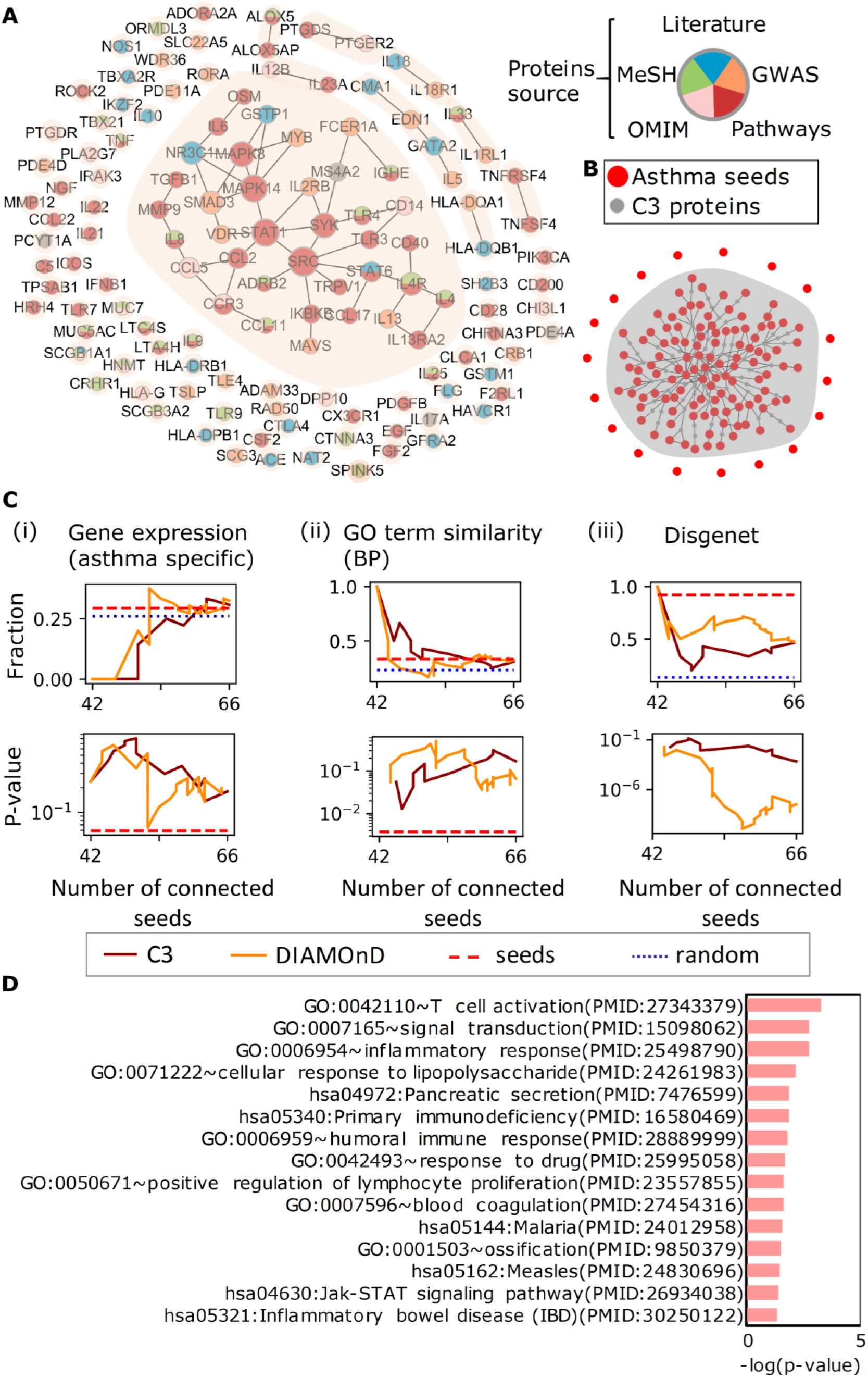
The visualization and biological explanation of asthma C3 module. (A) Subgraph of the full interactome showing the connections among the asthma DAPs. Asthma DAPs are scattered into 83 SCCs in the interactome (shown in shades). The colors of the nodes indicate the source of each seed. There are five sources: (i) Literature; (ii) GWAS; (iii) OMIM; (iv) MeSH; (v) Pathways. (B) Asthma C3 module. There are 111 seeds (red nodes) and 61 C3 proteins (grey nodes) in the module. The interaction between them is also indicated in the module. (C) Biological explanation of asthma C3 module. We consider three verification datasets: (i) Gene expression of asthma specific compiled from eight sources; (ii) GO term similarity (BP); (iii) Disgenet, number enrichment shows the fraction of the number of C3 and DIAMOnD proteins enriched on different validation datasets, statistical significance shows the corresponding enriched p-value which is derived from the hypergeometric distribution. (D) GO term (BP) and KEGG pathway functional annotation (p-value < 0.01) of C3 proteins used DAVID, the pink bar indicates the −log10 operation for the p-value obtained from the enrichment.

Furthermore, in order to better exploit the role of C3 proteins in the underlying disease pathways, we use DAVID^29^ (https://david.ncifcrf.gov/) to study the functional annotation of the 61 C3 proteins of asthma (Figure 3D). Here we focus on the GO terms (BP) and KEGG pathway function annotation. We find that C3 proteins are enriched on some pathways with a low p-value which has been reported to be associated with asthma (with PMID), such as: neuropeptide Y may enhance TH2 inflammatory response in asthma^30^, signal transduction of IL-13 plays role in the pathogenesis of bronchial asthma^31^, humoral immune responses during asthma^32^. And in table 2, we show in detail the enrichment function term and enriched C3 genes quantity and the PMID verified by the literature. Some C3 proteins are highly associated with asthma, such as: CD80 influences blockade on IL-4 and IFN-gamma production in nonatopic bronchial asthma^33^, variants in IL12A influence cockroach allergy among children with asthma^34^, ADA polymorphisms are related to asthma^35^. We demonstrate that the C3 module has biological significance. It can explain the disease mechanism and help predict potential pathogenic genes or novel drug target.

**Table 2.**
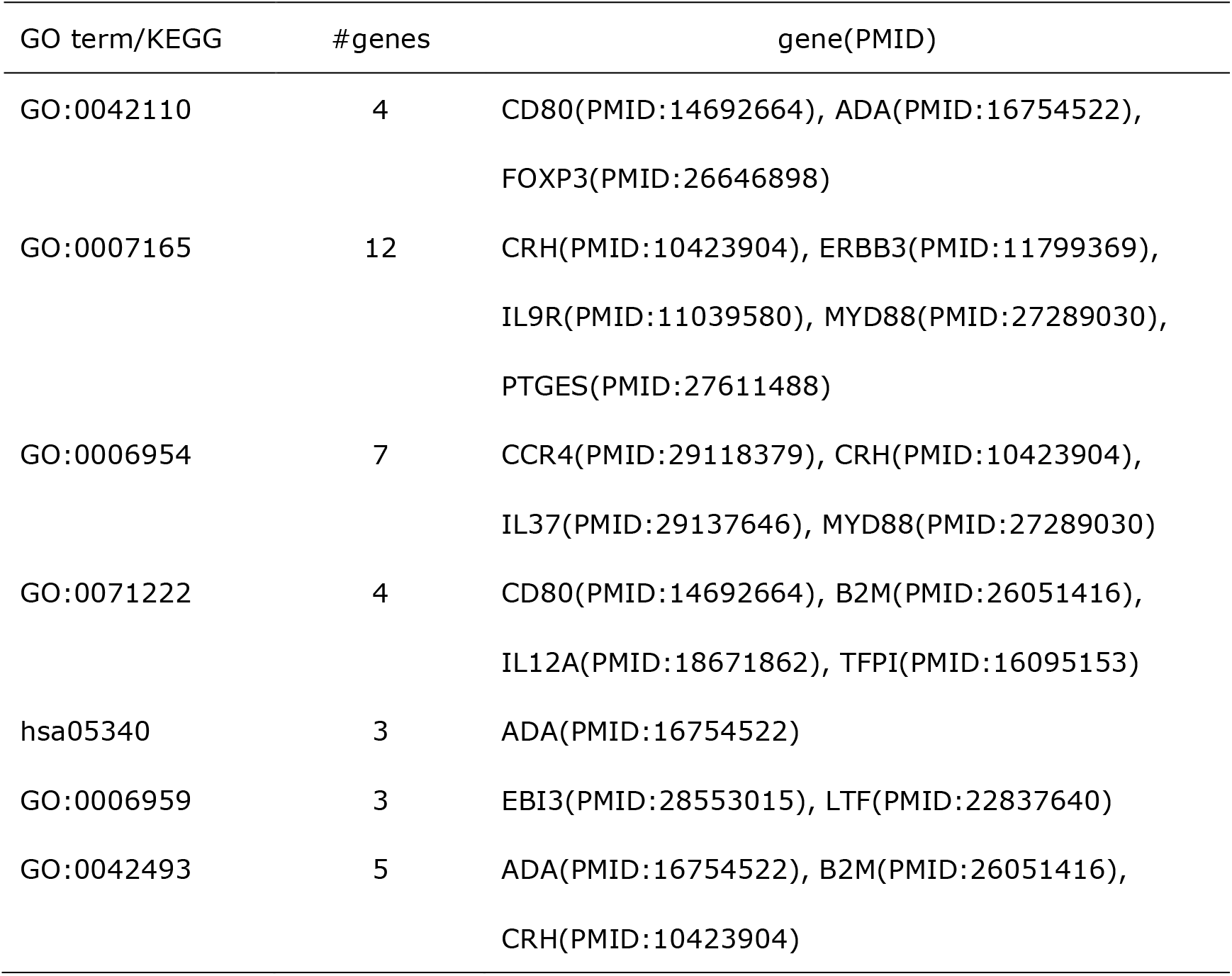
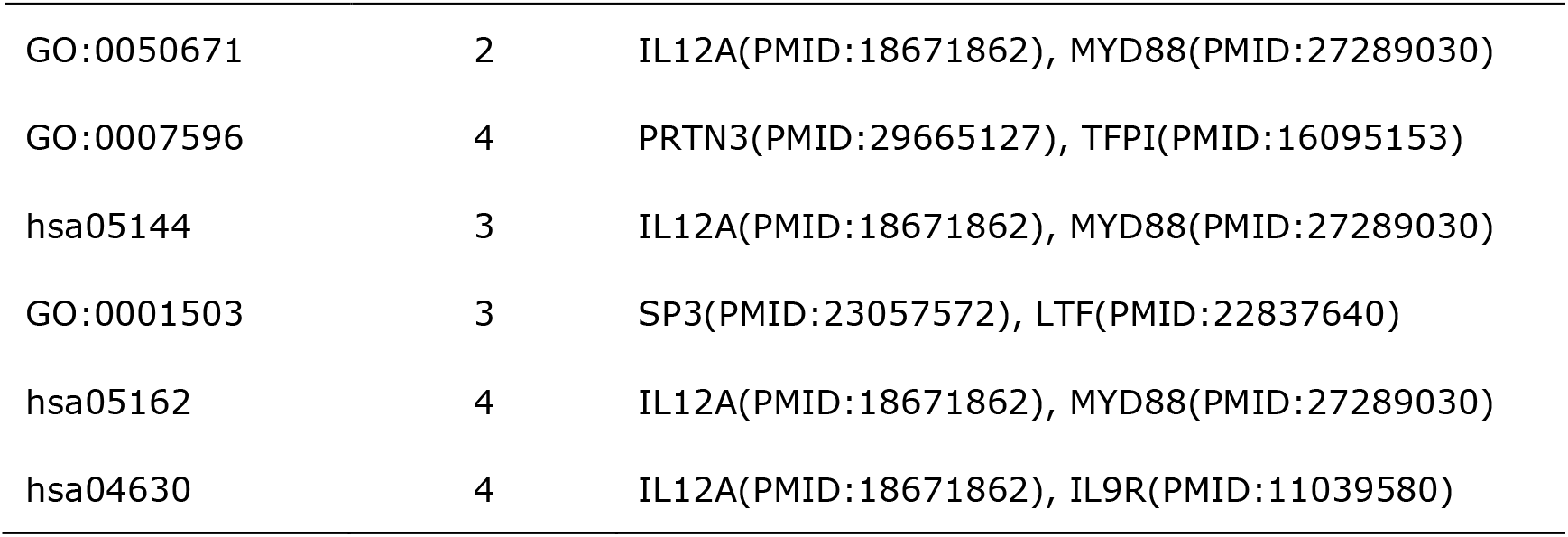
GO term and KEGG pathway functional annotation of C3 proteins and literature-validated PMID of C3 genes associated with asthma.

### C3 module in TCGA multi-omics data of BRCA

C3 method is not only used for DAPs, but also can be applied to the analysis at the whole genome level to obtain an omni genic^16^ disease neighbourhood. We collect four omics data: (i) mRNA expression; (ii) mutation; (iii) DNA methylation and (iv) copy number variation from TCGA (see Methods), and successfully identify multi-omics neighbourhoods of BRCA. In order to discover the connection between multi-omics data, we use the same human interactome network. For multi-omics data of BRCA, we identify the altered genes and take their encoding proteins as seeds, and import these seeds into C3 method to obtain their respective C3 module and C3 proteins (Figure 4A). Then, we compare different omics levels of mRNA expression, mutation, DNA methylation for C3 proteins, they are only significantly higher on their own omics levels. But in particular, for the C3 proteins of copy number variation, their mRNA expression and mutation levels are higher too (Figure 4B). For a same disease, is there any potential association between four omics data? We further want to see the relationship of their respective C3 proteins, we use the overlap to represent. As a result, we find that there is no significant association between four omics data, although all are related to a same disease. The same for their respective C3 proteins, there is no overlap between them, indicating that C3 proteins of the four omics data are also specific. That is, for BRCA, four specific omics data correspond to four specific groups of C3 proteins (Figure 4C).

**Figure 4.**
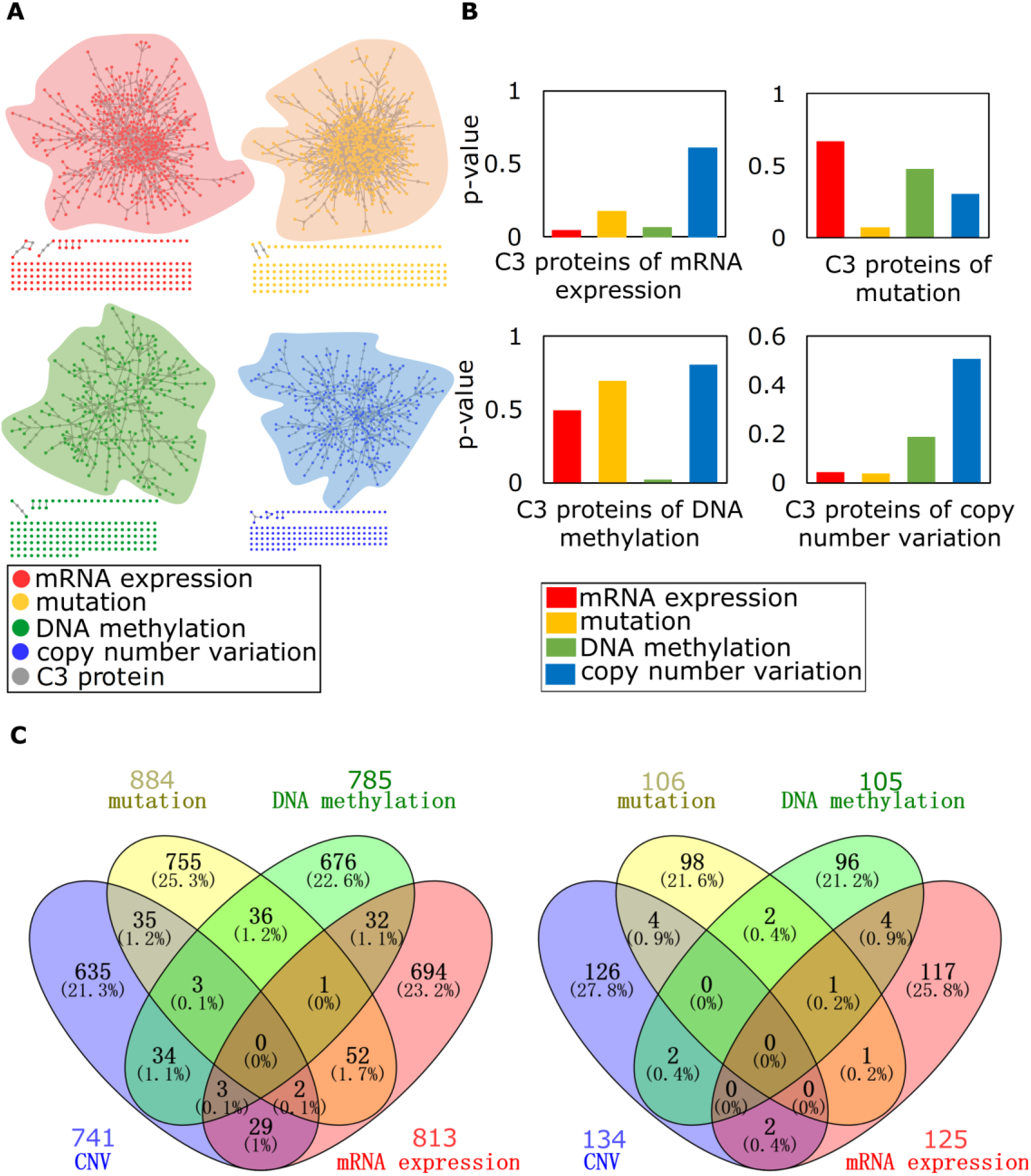
C3 module in TCGA multi-omics data of BRCA. (A) The C3 module of four omics data of BRCA: (i) mRNA expression; (ii) mutation; (iii) DNA methylation; (iV) copy number variation. (B) Validation of C3 proteins from four omics data. Red, yellow, green, and blue bars indicate level of four omics data compared to the rest of the network (p-value obtain by wilcoxon test). (C) Venn diagram of four omics data (left) and four different C3 proteins we obtained (right).

In table 3, we validate biological significance of four omics C3 proteins of BRCA, we consider three verification datasets: (i) OMIM; (ii) GWAS; (iii) Drug-target, for each validation dataset we use the wilcoxon test for enrichment. OMIM and GWAS databases can indicate the association between C3 proteins and diseases, and Drug-target database can observe that C3 proteins are associated with the drug target. We can find that the C3 proteins corresponding to these four omics data have different degrees of enrichment significance on these three verification datasets, which indicates that the C3 proteins recognized by our C3 method have different aspects, they are drug target, or in OMIM and GWAS.

**Table 3.** Biological significance of four omics C3 proteins of BRCA.

| Data<br>(#proteins) | mRNA<br>expression<br>(125)<br>#proteins (p-value) | mutation(106)<br>#proteins<br>(p-value) | DNA<br>methylation<br>(105)<br>#proteins (p-value) | CNV(134)<br>#proteins (p-value) |
| --- | --- | --- | --- | --- |
| OMIM(7952) | 100(**0**) | 75(**0.0056**) | 85(**0**) | 91(**0.01652**) |
| GWAS(5022) | 61(**0.00371**) | 57(**0.00022**) | 50(**0.01305**) | 53(0.2789) |
| Drug-target(2635) | 40(**0.00035**) | 32(**0.00328**) | 38(**0.00002**) | 34(**0.04129**) |

PPI network:13460 proteins

In order to further specifically assess the biological functions of the C3 proteins, we use ClueGO^36^ conduct GO term (BP) enrichment analysis (p-value < 0.01) of these C3 proteins (Figure 5). And enriched GO terms are indispensable pathway for the human such as: endocrine system development, cardiac muscle cell development, glucose transmembrane transport, positive regulation of collagen metabolic process. Many of them are also associated with BRCA, which again indicates it is feasible to apply C3 in multi-omics data. Our method can be used to identify different biologically active C3 proteins of different omics.

**Figure 5.**
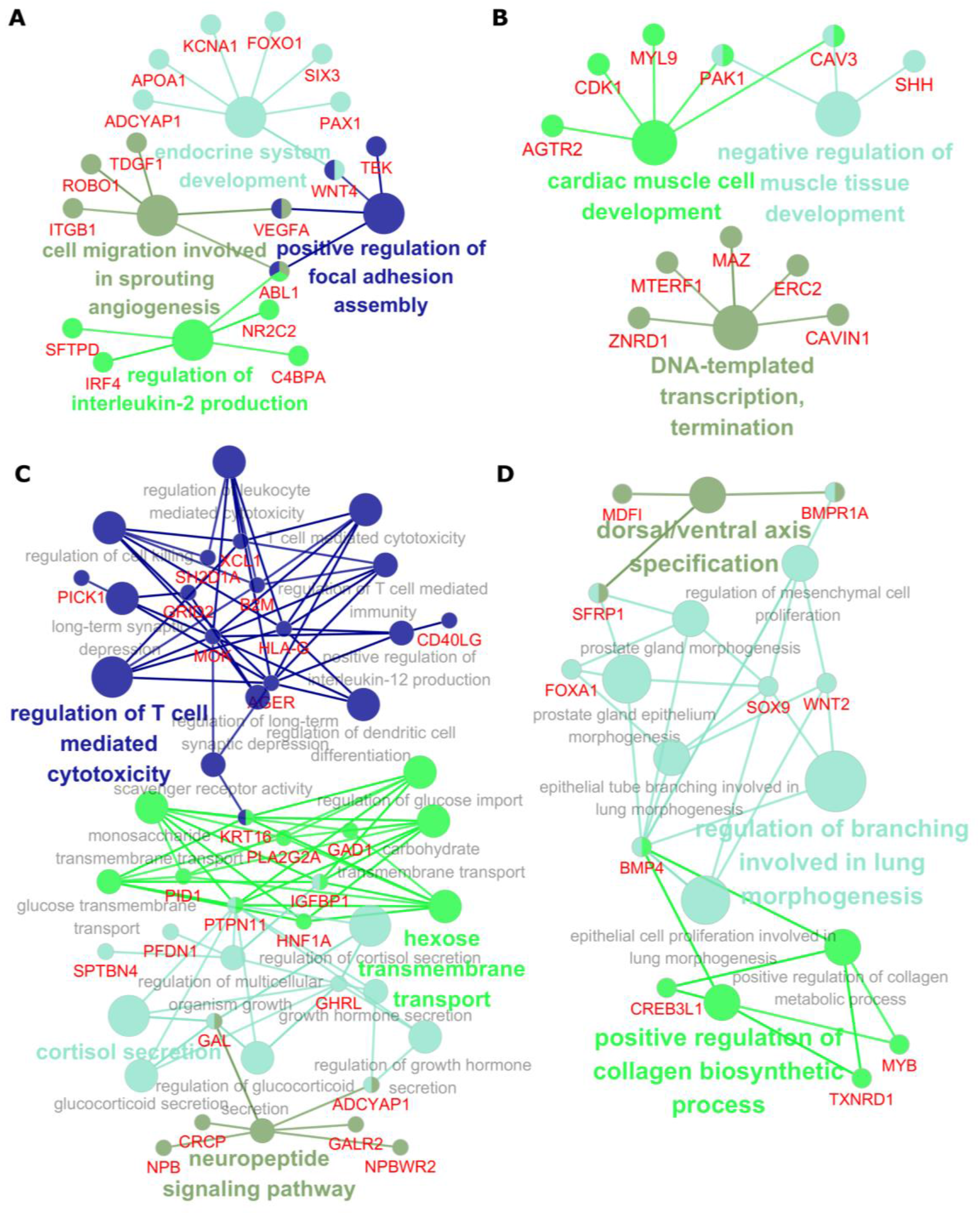
Biological significance of C3 proteins from four omics data of BRCA. (A,B,C,D) GO (BP) enrichment analysis of C3 proteins obtained from (A) mRNA expression, (B) mutation, (C) DNA methylation, and (D) copy number variation. Enrichment analysis is validated by tool ClueGO and the enriched proteins are represented by red nodes.

## Discussion

Disease module is helpful in understanding the molecular mechanism of disease causation and identifying novel drug target. Topological properties of disease module in the human interactome underlie all network-based prioritization tools for studying human diseases. However, DAPs appear to form plenty of SCCs in the incomplete interactome. The problem we solved is how to determine their connectivity pattern and detect a succinct disease module. We designed a C3 method by importing C3 proteins and C3 interactions with high connectivity to connect SCCs in the interactome. Our results highlight the role of C3 proteins and C3 interactions in connecting DAPs, which scattered as SCCs in incomplete interactome. Then we compared our method with DIAMOnD and random simulations to verify the succinctness and connectivity significance of the C3 module we obtained. As a matter of fact, C3 method can effectively construct succinct disease modules with relatively few C3 proteins for 299 disease from OMIM and GWAS databases and multi-omics data of BRCA from TCGA. And the disease module helps us to understand the disease pathways and identify pathogenic genes. We also have some unexpected results, there are no relationships between the different omics data for a same disease. Biologically, essential for disease characteristics, C3 proteins and C3 interactions may offer better and more accurate candidates. In the end, we wish our method can help to study potential disease mechanisms, disease heterogeneity, drug response, capture novel pathways and genes.

## Materials and methods

### Networks Information

For the accuracy of our experiment, we used the same network as DIAMOnD^9^: the human interactome. It comes from seven sources of protein interactions combined by Menche, J., et al: (i) regulatory interactions derived from transcription factors binding to regulatory elements; (ii) binary interactions from several yeast two-hybrid high-throughput and literature-curated data sets; (iii) literature-curated interactions derived mostly from low-throughput experiments;(iv) metabolic enzyme-coupled interactions; (v)protein complexes; (vi) kinase-substrate pairs; and (vii) signaling interactions. The union of all interactions from (i) to (vii) yielded a network of 13,460 proteins that were interconnected by 141,296 interactions^10^. In addition, we used three other networks to supplement the experiment: We took a current interactome from BioGRID (https://thebiogrid.org/) and used the latest release compiled on June, 2019 and only selected human interactions, finally we got 17,741 genes and 338,793 interactions. And in order to avoid the observed performance is not solely due to study biases in human interactome, we got networks from Bioplex (https://bioplex.hms.harvard.edu/) and HuRI (http://interactome.dfci.harvard.edu/H_sapiens/), finally Bioplex had 10,961 genes and 56,553 interactions and HuRI had 7,352 genes and 38,414 interactions. Network data was included into supplementary files.

### Disease-associated genes

Susan Dina Ghiassian et al.^15^ integrated DAPs from OMIM (Online Mendelian Inheritance in Man; http://www.ncbi.nlm.nih.gov/omim)^21^ and GWAS (Genome-Wide Association Studies. The DAPs from GWAS were obtained from the Phenotype-Genotype Integrator database (PheGenI; www.ncbi.nlm.nih.gov/gap/PheGenI)^8^ that integrates various NCBI genomic databases. They used a genome-wide significance cutoff of p-value ≤ 5.0e-8.Then in order to combine OMIM and GWAS they used the MeSH (Medical Subject Headings; http://www.nlm.nih.gov/mesh/) vocabulary. Using the hierarchical structure of the MeSH classification, in the result they found at least 20 DAPs and DAPs for which they have interaction information, and obtained 299 diseases and 3173 associated proteins.

### TCGA multi-omics data

We obtained publicly available TCGA (The Cancer Genome Atlas) multi-omics data of BRCA from UCSC Cancer Browser^37^ (https://xena.ucsc.edu). For mRNA expression, we used an R packet edge^38^ and took the threshold of |log(FC) | > 2 and adj.p-value < 0.001 to get 860 mRNA expression genes, where 632 mapped to human interactome were taken as DAPs. We used TCGA BRCA somatic mutation data and chose 889 genes that mutation counts > 10 and mutation frequency > 0.01, where 689 mapped were taken as DAPs. DNA methylation values, described as beta values, we used an R packet limma^39^ and choose 837 genes of the threshold |log(FC)| > 0.19 and adj.p-value < 1.0e^−14^, and only 367 mapped were taken as DAPs. BRCA thresholded gene-level copy number variation (CNV) estimated values to −2, −1, 0, 1, 2, representing homozygous deletion, single copy deletion, diploid normal copy, low-level copy number amplification, or high-level copy number amplification. Then we obtained 863 genes of threshold values −2 and 2 and only 446 mapped were taken as DAPs. After processing data, we obtained four altered data sets of multi-omics DAPs for BRCA, which are mRNA expression, mutation, DNA methylation and copy number variation.

### Biological validation datasets

Following data is provided in supplementary files.

(i) Gene expression of asthma specific compiled from eight sources: we selected eight expression data of direct asthma relevance from the Gene Expression Omnibus (GEO; https://www.ncbi.nlm.nih.gov/geo). We analyzed samples from GSE470, GSE473, GSE3004, GSE16032, GSE18965, GSE31773, GSE4302, GSE2125 with GEO2R, a total of 4111 differentially expressed genes were obtained where only 3273 in human interactome.

(ii) Gene ontology term similarity (BP): To elucidate the biological processes associated with the DAPs we used the GO biological process category. We calculated the GO term similarity (BP) of all genes, took the top 4000 proteins with high similarity to all DAPs, and only 3154 in human interactome. The calculation tool was R package GOSemSim^28^

(iii) Disgenet: Disgenet data of asthma got from DisGeNET database (http://www.disgenet.org/). Sources of data include literature data, inferred data, and animal models data, totally we got 1817 genes associate with the asthma.

(iv) OMIM: OMIM genes got from Online Mendelian Inheritance in Man (OMIM; https://omim.org), totally we got 9915 genes.

(v) GWAS: GWAS genes got from Genome-Wide Association Studies (GWAS; http://www.ebi.ac.uk/gwas), totally we got 15858 genes.

(vi) Drug-target: Drug-target genes obtained from The Drug Gene Interaction Database (DGIdb; http://www.dgidb.org), totally we got 2993 genes

### Connectivity of C3 protein and C3 interaction

The human interactome which can be denoted by network *G*(*N*, *E*), with proteins set *N* and edges set *E*. For a disease, we divide set *N* into set *D* of seeds and set *ND* of non-seeds, identify the SCCs of *D* in *G*, and construct the set *S* of all SCCs. We use *s*_0_ = |*S*| to indicate the number of SCCs needs to be connected, *n*_0_ = |*ND*| shows the number of non-seeds that can be the candidate proteins. For a protein *i*, its connectivity degree to SCCs (CDS) is defined as:

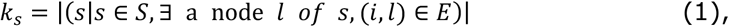

and its connectivity degree to non-seeds (CDN) is defined as:

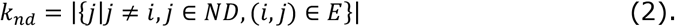

Overall, the connectivity degree of protein *i* calculate as *k* = *k*_*s*_ + *k*_*nd*_. Analogously, for an interaction *e*(*p*, *k*) between protein *p* and *k*, we combine the ability of two endpoints *p* and *k* together to quantify the CDS and CDN for *e*. Then, for *s_0_* randomly scatter SCCs, the probability that a protein or an interaction with a total connectivity degree *k* has exactly *ks* links to *s_0_* separate connected components is given by the hypergeometric distribution^40^:

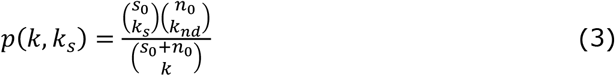

To evaluate whether a certain protein or an interaction has more connections to *s*_0_ SCCs than expected under this null hypothesis, we calculate the connectivity significant p-value, i.e. the cumulative probability for the observed or any higher number of connections:

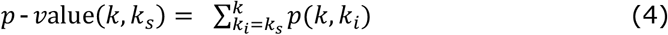

Then this p-value is used as a connectivity significant measure, and is use to quantify the abilities of protein and interaction in connecting separate connected components.

### C3 method

Suppose a network containing *s*_0_ SCCs formed by seeds of a particular disease and a relatively large number (*n*_0_) of non-seeds in the network. A greedy process can be used to select the most effective C3 proteins and C3 interactions to form the neighbourhood of a particular disease based on connectivity significance. Its upper bound complexity is *O*(*n*^2^).

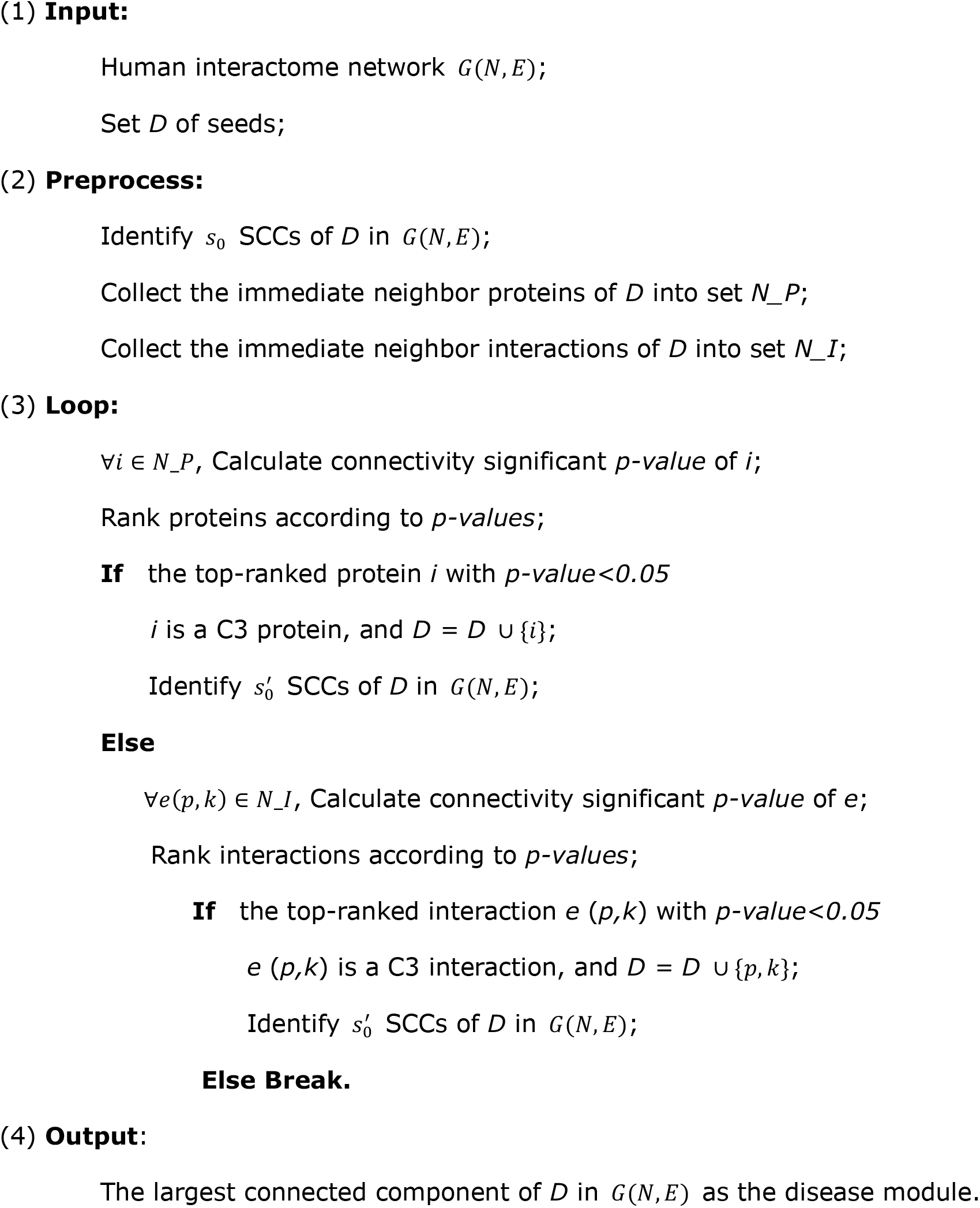

We implemented C3 with Python and users can download the code from https://github.com/wangbingbo2019/C3.

## Supporting Information

The experimental data: the networks data, disease genes from OMIM and GWAS, and some datasets for validation.

The C3 tool in Python.

The results of C3 modules.

## Acknowledgments

This work is supported by the National Natural Science Foundation of China (No. 61772395, 61432010, 61532014, 61672406, 61672407, 61702396 & 61702397), China Postdoctoral Science Foundation (No. 2015M582620), Fundamental Research Funds for the Central Universities (No. JB190306) and Shanghai Municipal Science and Technology Major Project (No.2018SHZDZX01), LCNBI and ZJLab.

## Author Contributions

BBW, JH conceived and designed the experiments. JH performed the experiments. CXZ, YJZ, LY, XLG, LG, YRC analyzed the data. BBW, JH, CXZ, YRC wrote the paper.

## Additional information

Competing financial interests: The authors declare no competing financial and non-financial interests.

## References

1. Goh, K.-I. et al. The human disease network. Proc. Natl. Acad. Sci. 104, 8685–8690 (2007).

2. Zanzoni, A., Soler-López, M. & Aloy, P. A network medicine approach to human disease. FEBS Lett. 583, 1759–1765 (2009).

3. Barabási, A.-L., Gulbahce, N. & Loscalzo, J. Network medicine: a network-based approach to human disease. Nat. Rev. Genet. 12, 56–68 (2011).

4. Schadt, E. E. Molecular networks as sensors and drivers of common human diseases. Nature 461, 218–223 (2009).

5. Pawson, T. & Linding, R. Network medicine. FEBS Lett. 582, 1266–1270 (2008).

6. Califano, A., Butte, A. J., Friend, S., Ideker, T. & Schadt, E. Leveraging models of cell regulation and GWAS data in integrative network-based association studies. Nat. Genet. 44, 841–847 (2012).

7. Venkatesan, K. et al. An empirical framework for binary interactome mapping. Nat. Methods 6, 83–90 (2009).

8. Ramos, E. M. et al. Phenotype–Genotype Integrator (PheGenI): synthesizing genome-wide association study (GWAS) data with existing genomic resources. Eur. J. Hum. Genet. 22, 144–147 (2014).

9. Ghiassian, S. D., Menche, J. & Barabási, A.-L. A DIseAse MOdule Detection (DIAMOnD) Algorithm Derived from a Systematic Analysis of Connectivity Patterns of Disease Proteins in the Human Interactome. PLOS Comput Biol 11, e1004120 (2015).

10. Menche, J. et al. Uncovering disease-disease relationships through the incomplete interactome. Science 347, 1257601 (2015).

11. Agrawal, M., Zitnik, M. & Leskovec, J. Large-Scale Analysis of Disease Pathways in the Human Interactome. bioRxiv 189787 (2017). doi:10.1101/189787

12. Vinayagam, A. et al. Controllability analysis of the directed human protein interaction network identifies disease genes and drug targets. Proc. Natl. Acad. Sci. U. S. A. 113, 4976–4981 (2016).

13. Ghiassian, S. D. et al. Endophenotype Network Models: Common Core of Complex Diseases. Sci. Rep. 6, 27414 (2016).

14. Huttlin, E. L. et al. Architecture of the human interactome defines protein communities and disease networks. Nature 545, 505–509 (2017).

15. Liu, C.-C. et al. DiseaseConnect: a comprehensive web server for mechanism-based disease-disease connections. Nucleic Acids Res. 42, W137–146 (2014).

16. Boyle, E. A., Li, Y. I. & Pritchard, J. K. An Expanded View of Complex Traits: From Polygenic to Omnigenic. Cell 169, 1177–1186 (2017).

17. Wray, N. R., Wijmenga, C., Sullivan, P. F., Yang, J. & Visscher, P. M. Common Disease Is More Complex Than Implied by the Core Gene Omnigenic Model. Cell 173, 1573–1580 (2018).

18. Sharma, A. et al. A disease module in the interactome explains disease heterogeneity, drug response and captures novel pathways and genes in asthma. Hum. Mol. Genet. 24, 3005–3020 (2015).

19. Navlakha, S. & Kingsford, C. The power of protein interaction networks for associating genes with diseases. Bioinformatics 26, 1057–1063 (2010).

20. Radicchi, F., Castellano, C., Cecconi, F., Loreto, V. & Parisi, D. Defining and identifying communities in networks. Proc. Natl. Acad. Sci. 101, 2658–2663 (2004).

21. Hamosh, A., Scott, A. F., Amberger, J. S., Bocchini, C. A. & McKusick, V. A. Online Mendelian Inheritance in Man (OMIM), a knowledgebase of human genes and genetic disorders. Nucleic Acids Res. 33, D514–D517 (2005).

22. Welter, D. et al. The NHGRI GWAS Catalog, a curated resource of SNP-trait associations. Nucleic Acids Res. 42, D1001–1006 (2014).

23. Sadeghi, A. & Fröhlich, H. Steiner tree methods for optimal sub-network identification: an empirical study. BMC Bioinformatics 14, 144 (2013).

24. Tumminello, M., Miccichè, S., Lillo, F., Piilo, J. & Mantegna, R. N. Statistically Validated Networks in Bipartite Complex Systems. PLOS ONE 6, e17994 (2011).

25. Callaway, D. S., Newman, M. E., Strogatz, S. H. & Watts, D. J. Network robustness and fragility: percolation on random graphs. Phys. Rev. Lett. 85, 5468–5471 (2000).

26. Sanchez-Vega, F. et al. Oncogenic Signaling Pathways in The Cancer Genome Atlas. Cell 173, 321–337.e10 (2018).

27. Geoffrey J. McLachlan, Kim‐Anh Do & Christophe Ambroise. Analyzing Microarray Gene Expression Data. Wiley. (2004).

28. Yu, G. et al. GOSemSim: an R package for measuring semantic similarity among GO terms and gene products. Bioinforma. Oxf. Engl. 26, 976–978 (2010).

29. Huang, D. W. et al. DAVID Bioinformatics Resources: expanded annotation database and novel algorithms to better extract biology from large gene lists. Nucleic Acids Res. 35, W169–W175 (2007).

30. Lu, Y. et al. Neuropeptide Y may mediate psychological stress and enhance TH2 inflammatory response in asthma. J. Allergy Clin. Immunol. 135, 1061–1063.e4 (2015).

31. Izuhara, K. & Arima, K. Signal transduction of IL-13 and its role in the pathogenesis of bronchial asthma. Drug News Perspect. 17, 91–98 (2004).

32. Doorley, L. A., LeMessurier, K. S., Iverson, A. R., Palipane, M. & Samarasinghe, A. E. Humoral immune responses during asthma and influenza co-morbidity in mice. Immunobiology 222, 1064–1073 (2017).

33. Rutkowski, R. et al. CD80 and CD86 expression on LPS-stimulated monocytes and the effect of CD80 and CD86 blockade on IL-4 and IFN-gamma production in nonatopic bronchial asthma. Arch. Immunol. Ther. Exp. (Warsz.) 51, 421–428 (2003).

34. Pistiner, M. et al. Polymorphisms in IL12A and cockroach allergy in children with asthma. Clin. Mol. Allergy CMA 6, 6 (2008).

35. Liu, Y. et al. ADA polymorphisms and asthma: a study in the Chinese Han population. J. Asthma Off. J. Assoc. Care Asthma 43, 203–206 (2006).

36. Bindea, G. et al. ClueGO: a Cytoscape plug-in to decipher functionally grouped gene ontology and pathway annotation networks. Bioinforma. Oxf. Engl. 25, 1091–1093 (2009).

37. Mary Goldman, Brian Craft, Mim Hastie, Kristupas Repečka, Akhil Kamath, Fran McDade, Dave Rogers, View ORCID ProfileAngela N Brooks, Jingchun Zhu, David Haussler. The UCSC Xena Platform for cancer genomics data visualization and interpretation | bioRxiv. Available at: https://www.biorxiv.org/content/10.1101/326470v4. (Accessed: 3rd April 2019)

38. Leek, J. T., Monsen, E., Dabney, A. R. & Storey, J. D. EDGE: extraction and analysis of differential gene expression. Bioinforma. Oxf. Engl. 22, 507–508 (2006).

39. Ritchie, M. E. et al. limma powers differential expression analyses for RNA-sequencing and microarray studies. Nucleic Acids Res. 43, e47 (2015).

40. John A, R. Mathematical Statistics and Data Analysis (Third ed.).Duxbury Press. (2007).

